# Inflammation-mediated Upregulation of VCAM-1 but not KIM-1 during Acute Kidney Injury to Chronic Kidney Disease Transition

**DOI:** 10.1101/2022.09.15.508151

**Authors:** Isabel Melchinger, Kailin Guo, Jiankan Guo, Leyuan Xu

## Abstract

**Background:** Patients with acute kidney injury (AKI) have higher risks of developing chronic kidney disease (CKD). The basis for the AKI-to-CKD transition remains poorly understood, but studies in animal models suggest a linkage between the inflammatory response to injury and subsequent nephron loss and interstitial fibrosis. The proximal tubule is the primary venue of injury and progression of disease during this process.

**Methods:** Mouse unilateral ischemia/reperfusion injury (U-IRI) model was used to study the kinetics of proximal injury marker expression during AKI-to-CKD transition. Immortalized MPT cells and primary cultured renal cells were used to study factor(s) that induce vascular cell adhesion protein-1 (VCAM-1) expression in proximal tubule cells.

**Results:** Kidney injury molecule-1 (KIM-1) was rapidly upregulated on day 1 after injury and gradually reduced close to the baseline; whereas VCAM-1 was not upregulated on day 1 but markedly increased afterwards during AKI-to-CKD transition. The proximal tubular VCAM-1 expression is induced by proinflammatory cytokines including TNFα and IL-1β. Blockade of these signaling pathways by using NF-κB inhibitor or by using double null mutant *Myd88* and *Trif* derived PCRC in vitro or decrease of immune cell recruitment using *Ccr2* null mouse in vivo significantly suppressed VCAM-1 expression. Human single cell transcriptome analysis identified a distinct cluster of injured proximal tubules that highly expressed *VCAM1* but not *HAVCR1 (KIM1*), and the population of these VCAM1-positive proximal tubule cells was associated with CKD progression and VCAM-1 levels were significantly higher in the patients with Stage 3 CKD as compared to the healthy references.

**Conclusions:** Proximal tubule cells upregulated KIM-1 and VCAM-1 in an orchestrated fashion after injury. Upregulation of VCAM-1 associated with chronic tubular injury and interstitial fibrosis and may mark the earliest molecular event during AKI-to-CKD transition.

## INTRODUCTION

Patients with acute kidney injury (AKI) have higher risks for developing chronic kidney disease (CKD) and end-stage renal disease (ESRD) (1). The basis for the AKI-to-CKD transition remains poorly understood, but studies in animal models suggest a linkage between the inflammatory response to injury and subsequent nephron loss and interstitial fibrosis (2). The proximal tubule becomes the primary target of injury and progression of kidney disease (3). Following ischemia/reperfusion injury (IRI), the proximal tubules especially S3 segment and thick ascending limbs undergo necroptosis via cell membrane and possibly nuclear envelope rapture and/or ferroptosis via lipid peroxidation, leading to a great loss of tubular cells (4, 5). In response to the initial injury, proximal tubule cells upregulate kidney injury molecule-1 (KIM-1), which acts as a phosphatidylserine phagocytosis and scavenger receptor that can bind to lipids on the surface of apoptotic and necrotic cells as well as to oxidized LDL to protect the kidney from further damage (6). Renal recovery after IRI requires both the replacement of those dead cells and the timely clearance of casts formed by the dead cell debris. Immune cells especially macrophages play an important role in promoting tubule repair following IRI (7–10). In cases of more severe injury (longer clamp times in IRI models or injury to older mice) or sustained injury (unilateral ureteral obstruction), macrophages remain in the kidney interstitium adjacent to nonrepaired tubules. Their activation is responsible for recruiting a second wave of immune cell infiltration into the unresolved kidneys. The persistence of interstitial immune cells in turn promotes kidney fibrosis and inflammation, which lead to CKD transition (11–15).

In this study, we identified two distinct expression kinetics of proximal tubule injury markers, KIM-1 and vascular cell adhesion protein-1 (VCAM-1), following unilateral ischemia/reperfusion injury (U-IRI). KIM-1 was rapidly upregulated on day 1 after injury and gradually reduced close to the baseline; whereas VCAM-1 was not upregulated on day 1 but markedly increased afterwards, suggesting the kidney undergoes two distinct injury phases during AKI-to-CKD transition. Using immortalized proximal tubule cell line and primary cultured renal cells (PCRCs), we demonstrated that the proximal tubularVCAM-1 expression is induced by proinflammatory cytokines including TNFα and IL-1β. Blockade of TNFα and IL-1β signaling pathways by using NF-κB inhibitor or knockout of *Myd88* and *Trif* in vitro or decrease of immune cell recruitment using *Ccr2* null mouse in vivo significantly suppressed VCAM-1 expression. Consistent with in vitro and in vivo models, using human single cell transcriptome analysis, we identified a distinct cluster of injured proximal tubules that highly expressed *VCAM1* but not *HAVCR1 (KIM1*), and the population of these VCAM1-positive proximal tubule cells was associated with CKD progression and significantly more abundant and in the patients with Stage 3 CKD as compared to the healthy references. Together, our results suggest that different from AKI, proximal tubule cells upregulated VCAM-1 but not KIM-1 in response to the inflammatory milieu, which was associated with kidney atrophy and kidney fibrosis.

## Methods

### Animal Surgery and Experimental Protocol

All animal protocols were approved by the Yale University Institutional Animal Care and Use Committee. C57BL/6 wild-type mice (Envigo) as well as *Ccr2*^−/−^ (Jackson Laboratory) and the suggested genetic background control C57BL/6J wild-type mice (Jackson Laboratory) (age 9-11 weeks) were used in this work. All mice were maintained on a 12-hour light and 12-hour dark cycle with free access to standard food and water before and after surgery. Due to the substantial difference in susceptibility to IRI injury between male and female mice (16), male mice were exclusively used to reduce total numbers of mice required for statistical analysis. Before surgery, all mice were subjected to anesthesia by intraperitoneal injection with ketamine (100 mg/kg) and xylazine (10 mg/kg) on a 37 °C warming pad. To establish the unilateral ischemia/reperfusion injury (U-IRI) model, the abdomen was opened, and warm renal ischemia was induced using a nontraumatic microaneurysm clip (FST Micro Clamps) on the left renal pedicle for 27 min, leaving the right kidney intact. During surgery, all mice received intraperitoneal phosphate-buffered saline (PBS) and buprenorphine (0.1 mg/kg) to avoid dehydration and postoperative pain, respectively. The mice were sacrificed on day 1, 14 and 30 after U-IRI (n=10 mice/end point). Baseline control mice were sacrificed and denoted as day 0 for the injury (n=10 mice) (15).

### In Vitro Cell Culture

#### MPT Cell Culture

Pathogen-free MPT cells (BU-MPT immortalized cells) were plated in 6-well plates in Dulbecco’s Modified Eagle Medium (DMEM) (Gibco) supplied with 10% FBS (Sigma-Aldrich) and Antibiotic-Antimycotic (Gibco) for 24 hrs and then treated with MPT cell debris, 500 μM H_2_O_2_, 10 ng/mL LPS (Sigma-Aldrich), 20 ng/mL TNFα (R&D Systems), 100 ng/mL interferon γ (R&D Systems), or serum depletion for 6 hrs. In a separated experiment, MPT cells were treated with 20 ng/mL TNFα (R&D Systems) or 20 ng/mL IL-1β (R&D Systems) in the presence or absence of a selective IKKα and IKKβ inhibitor, ACHP (R&D Systems) at 1 μM concentration for 6 hrs. After treatment, the cell lysates were harvested with RLT buffer (Qiagen) supplied with β-mercaptoethanol for RNA extraction.

#### Primary Cultured Renal Cell (PCRC) Culture

PCRCs were isolated from *Wild-type* or *Myd88^−/−^;Trif^−/−^* mice using our recently reported protocol with minor modification (10). Briefly, donor mouse kidney was minced in 2 mm^3^ cubes with a razor blade, and incubated with 5 mL Liberase (0.5 mg/mL, Roche) per kidney, supplemented with DNaseI (100 μg/mL, Roche) and MgCl_2_ (0.1%) in PBS for 30 min with gentle pipetting every 10 min. The digested mixture was passed through a 70 μm cell strainer (Falcon) mounted on a 50 mL tube, rinsed with 45 mL ice-cold PBS, and spun down at 1000 rpm for 5 min at 4 °C. The resuspended cells/tubule segments pellets were treated with 3 mL red blood cell lysis buffer (Alfa Aesar) at room temperature for 3 min to remove red blood cells. Primary culture was established in a complete medium: DMEM supplied with 10% FBS and Antibiotic-Antimycotic. After one medium change 48 hrs after seeding, the culture was continued for another 3 days, and PCRC were trypsinized and plated in 6-well plates (for RNA extraction) or 60-mm dishes (for protein extraction) in the complete media for 24 hrs and then treated with 20 ng/mL TNFα or 20 ng/mL IL-1β in the presence or absence of 1 μM ACHP for 6 hrs (for RNA extraction) or 24 hrs (for protein extraction). After treatment, the cell lysates were harvested with RLT buffer (Qiagen) supplied with β-mercaptoethanol for RNA extraction or RIPA buffer (Thermo Fisher Scientific) supplied with protease and phosphatase inhibitors (Thermo Fisher Scientific) for protein extraction.

### Quantitative PCR Analysis

Whole kidney RNA was extracted with an RNeasy Mini kit (Qiagen) and reverse transcribed using the iScript cDNA Synthesis Kit (Bio-Rad). Gene expression analysis was determined by quantitative real-time PCR using an iCycler iQ (Bio-Rad) and normalized to *Hprt*. The primers include *Hprt* forward: CAGTACAGCCCCAAAATGGT, reverse: CAAGGGCATATCCAACAACA; *Havcr1* forward: GAGAGTGACAGTGGTCTGTATTG, reverse: CGTGTGGGAATCTCTGGTTTA; and *Vcam1* forward: ACTCCCGTCATTGAGGATATTG, reverse: GTTGTATTCCTGGGAGAGATGTAG. The data were expressed using the comparative threshold cycle (ΔCT) method, and the mRNA ratios were given by 2^−ΔCT^.

### Immunofluorescence (IF)

Kidneys were fixed in 10% neutral buffered formalin and embedded in paraffin. Deparaffinized kidney sections were rehydrated in graded alcohols (100%, 95%, 90%, 80%, and 70%) and microwaved in citrate buffer antigen retrieval for 20 minutes. Sections were washed with PBS, blocked with 10% normal donkey serum, and then stained with primary antibodies against VCAM-1 (#32653, Cell Signaling Technology, 1:100 dilution) and KIM-1(#AF1817, Novus Biologicals, 1:100 dilution)(17). The sections were mounted with VECTASHIELD^®^ HardSet™ Antifade Mounting Medium with DAPI (4,6-diamidino-2-phenylindole). The fluorescence images were obtained by confocal microscopy (Zeiss LSM 880).

### Western Blot Analysis

Kidney lysates were fractioned using a RIPA lysis and extraction buffer (Thermo Fisher Scientific) supplied with protease and phosphatase inhibitors (Thermo Fisher Scientific). Protein concentration was measured using Bio-Rad Protein Assay. Fifty microgram of protein was separated by SDS electrophoresis using 10% polyacrylamide gel and transferred to an Immobilon PVDF membrane (Millipore). Membrane was blocked with 5% non-fat milk in TBST for 2 hrs and probed overnight at 4°C with the primary antibody against TIM-1/KIM-1/HAVCR (#AF1817, Novus Biologicals). After washing the membrane with TBST, it was incubated for 1 hour at room temperature with HRP-conjugated secondary antibody (Thermo Fisher Scientific). The membrane was developed using the ECL Detection System (Thermo Fisher Scientific) and imaged using the Odyssey Fc Imaging System (LI-COR Biosciences). Next, the membrane was washed with TBS and then immersed and incubated in Restore™ PLUS Western Blot Stripping Buffer (Thermo Fisher Scientific) for 15 min. After TBST wash, the same membrane was blocked with 5% non-fat milk in TBST for 2 hrs, probed overnight at 4°C with the primary antibody against VCAM-1 (#32653, Cell Signaling Technology), and developed using the ECL Detection System following the same protocol as described above. Lastly, the same membrane was stripped, washed, blocked, probed overnight at 4°C with the primary antibody against β-actin (AC-15) (#sc-69879, Santa Cruz Biotechnology), and developed using the ECL Detection System following the same protocol as described above. Western blot images were quantified using LICOR’s Image Studio Lite. Equivalently sized rectangles were placed around each band and sized to fully contain the largest band present. Background was subtracted using the median intensity value of a region extending 3 pixels beyond the boundary of the rectangle, from the top and bottom. Relative signal was reported as the total intensity, reduced by the product of the background and the area of the rectangle.

### Proteome Profiler Cytokine Array

Three *wild-type* or *Ccr2^−/−^* kidney lysates were equally pooled (60 μg/kidney, 180 μg/mL in final concentration for sample, i.e., *wild-type* and *Ccr2^−/−^*). Cytokine array was carried out using Mouse XL Cytokine Array Kit (R&D Systems, #ARY028) according to the manufacturer’s instructions.

### scRNA-seq Analysis of Mouse Kidneys

The scRNA-seq data library was generated from mice kidneys following U-IRI on days 0 (control), 7, 14, and 30 (n=2/time point). The library generation, data preprocessing and analysis were previously described (15). The gene expressions of proximal tubule injury markers were visualized across each time point.

### scRNA-seq Data Analysis of Human Kidneys

To generate the human scRNA-seq library, we accessed the scRNA-seq expression matrices and the corresponding clinical dataset from the publicly available Kidney Precision Medicine Project (KPMP) Central Biorepository (https://atlas.kpmp.org/). Downstream data analysis was performed using the Seurat v4.0 R package. In the initial quality control (QC) analysis, samples without mitochondria genes were excluded, and in the reference, the only transplant pre-perfusion biopsy samples were included as healthy controls, which results 10 patients with CKD and 8 healthy participants. The clinical datasets were summarized in the Table 1 and 2 (https://atlas.kpmp.org/). All the 18 objects were merged for additional QC analysis. Poor quality cells with <300 unique genes or <500 unique molecular identifier (UMI) counts (likely cell fragment) and >6,000 unique genes or >40,000 UMI (potentially cell duplet) were excluded. Cells were excluded if their mitochondrial gene percentages were over 50%. Low-complexity cells like red blood cells with <0.8 log10 genes per UMI counts were also excluded (18). Only genes expressed in 5 or more cells were used for further analysis. The QC filters resulted in a total of 58,357 cells with a median of 1,836 UMI counts per cell at a sequencing depth of 58,301 genes across 58,357 cells. Confounding sources of variation including mitochondrial gene content were removed for downstream clustering analysis. The merged dataset was split, normalized, cell cycle scored, SCTransformed, and integrated using 3,000 most variable genes (19–21). Principle component analysis (PCA) was performed on the integrated data. The top 20 principal components were chosen for cell clustering and neighbors finding with k.param = 20, perplexity of 30, and resolution = 0.7. The Uniform Manifold Approximation and Projection (UMAP) was used to visualize the single cells in two-dimensional space. Each cluster was screened for marker genes by differential expression analysis based on the non-parametric Wilcoxon rank sum test for all clusters with genes expressed in at least 25% of cells either inside or outside of a cluster. Based on the kidney cell and immune cell lineage-specific marker expression, 20 cell clusters were identified. The gene expressions of proximal tubule injury markers were visualized across all types of participants.

**Table 1.**
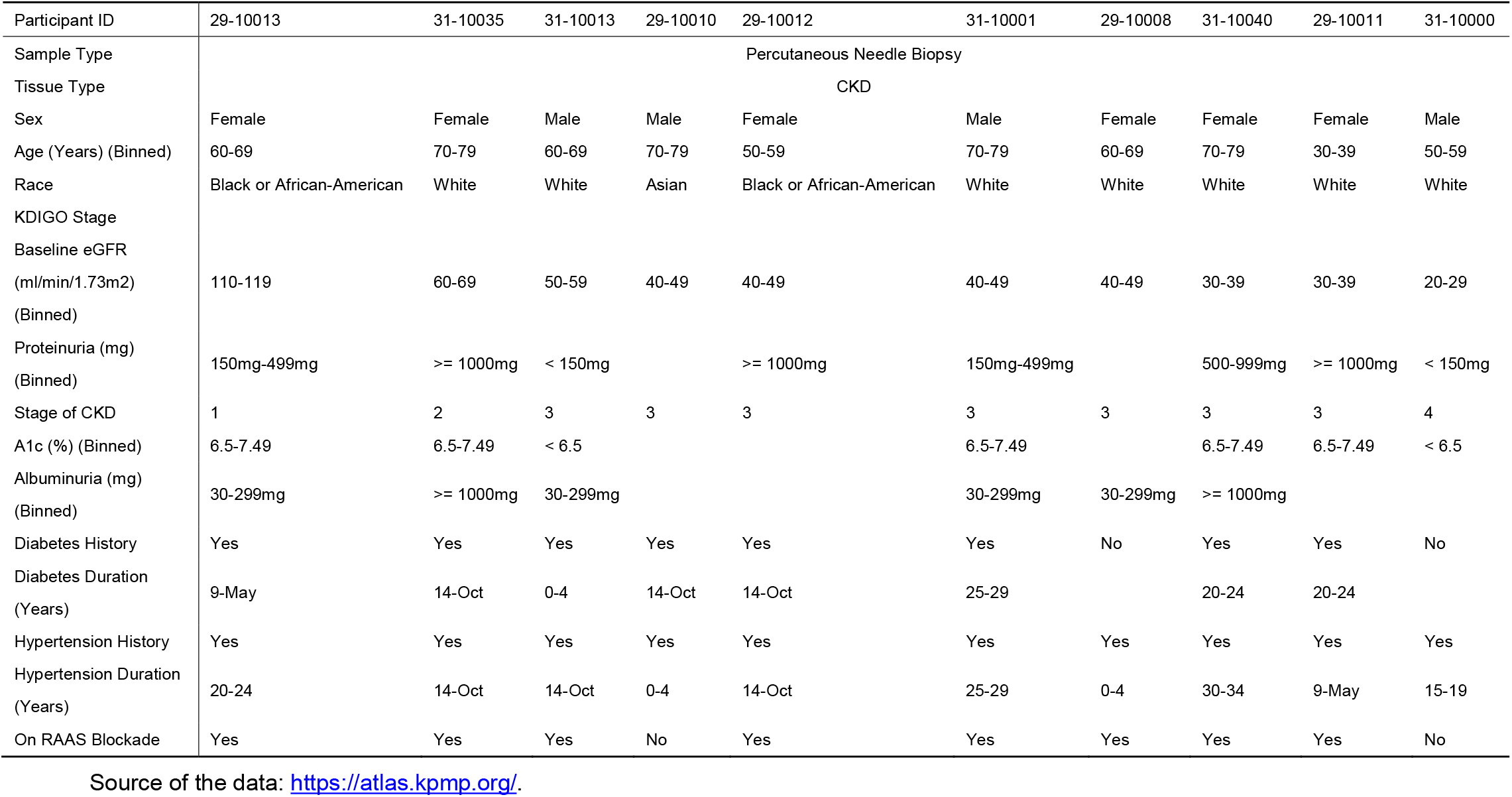
Clinical dataset of participants with CKD from the publicly available KPMP Central Biorepository.

**Table 2.**
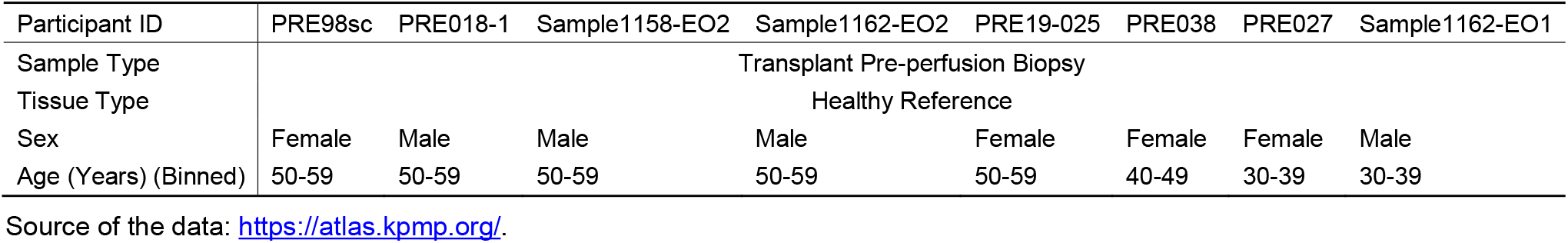
Clinical dataset of healthy participants from the publicly available KPMP Central Biorepository.

### Statistical Analysis

The data were expressed as means ± standard deviation (SD). Two-group comparison was performed by two-tailed Student’s *t*-test. Multigroup comparison was performed by one-way analysis of variance (ANOVA) for group mean comparison followed by Tukey’s multiple comparison test for subgroup comparison. Correlation of gene expression was performed by Pearson correlation coefficient R with two-tailed *P* value. All the statistical analysis was performed using Prism 8 (GraphPad Software). A value of *P*<0.05 was considered statistically significant.

## Results

### VCAM-1 but not KIM-1 is upregulated during AKI-to-CKD transition

We and others have shown that following U-IRI, the injured kidney fails to repair and instead atrophies and develops fibrosis at the late stage of IRI (11, 12, 14). Histologically, the atrophied kidney loses proximal tubular brush border along with a marked decrease of proximal tubule differentiation marker expression including megalin (*Lrp2*), NaPi-IIa (*Scl34a1*), and NaDC3 (*Slc13a3*) (15). Analysis of kidney section and whole kidney protein showed that the expression of proximal tubule injury marker KIM-1 was significantly increased on day 1, sustained on day 14, but significantly decreased on day 30 after IRI (Fig. 1A-C and Supplementary Figure 1). In contrast, VCAM-1 was not upregulated on day 1 but markedly increased afterwards up to day 30 after IRI and predominantly expressed in the proximal tubule epithelial cells (Fig. 1A-C and Supplementary Figure 1), consistent with our previous findings at the mRNA level (15). At the single cell transcriptome level (15), both *Vcam1* and *Kim1* (*Havcr1*) were highly upregulated by the injured proximal tubule epithelial cells on day 7 after IRI but diverged afterwards: downregulation of *Kim1* but sustained expression of *Vcam1* on day 14 and 30 after IRI (Figure 1D), consistent with the findings using bilateral IRI up to 6 weeks after IRI (22). The proximal tubules that express Vcam1 are proinflammatory and have regulon activity for both Relb and NF-κB, leading to failure of tubular repair (22). The whole kidney mRNA expression level of *Vcam1* but not the *Kim1* (*Havcr1*) highly correlated with those of fibrosis markers including *Col1a1, Col3a1, Fn1*, and *Pdgfrb* in the period of progressive atrophy seen in this model (Table 3), suggesting that VCAM-1 but not KIM-1 is associated with AKI-to-CKD transition.

**Figure 1.**
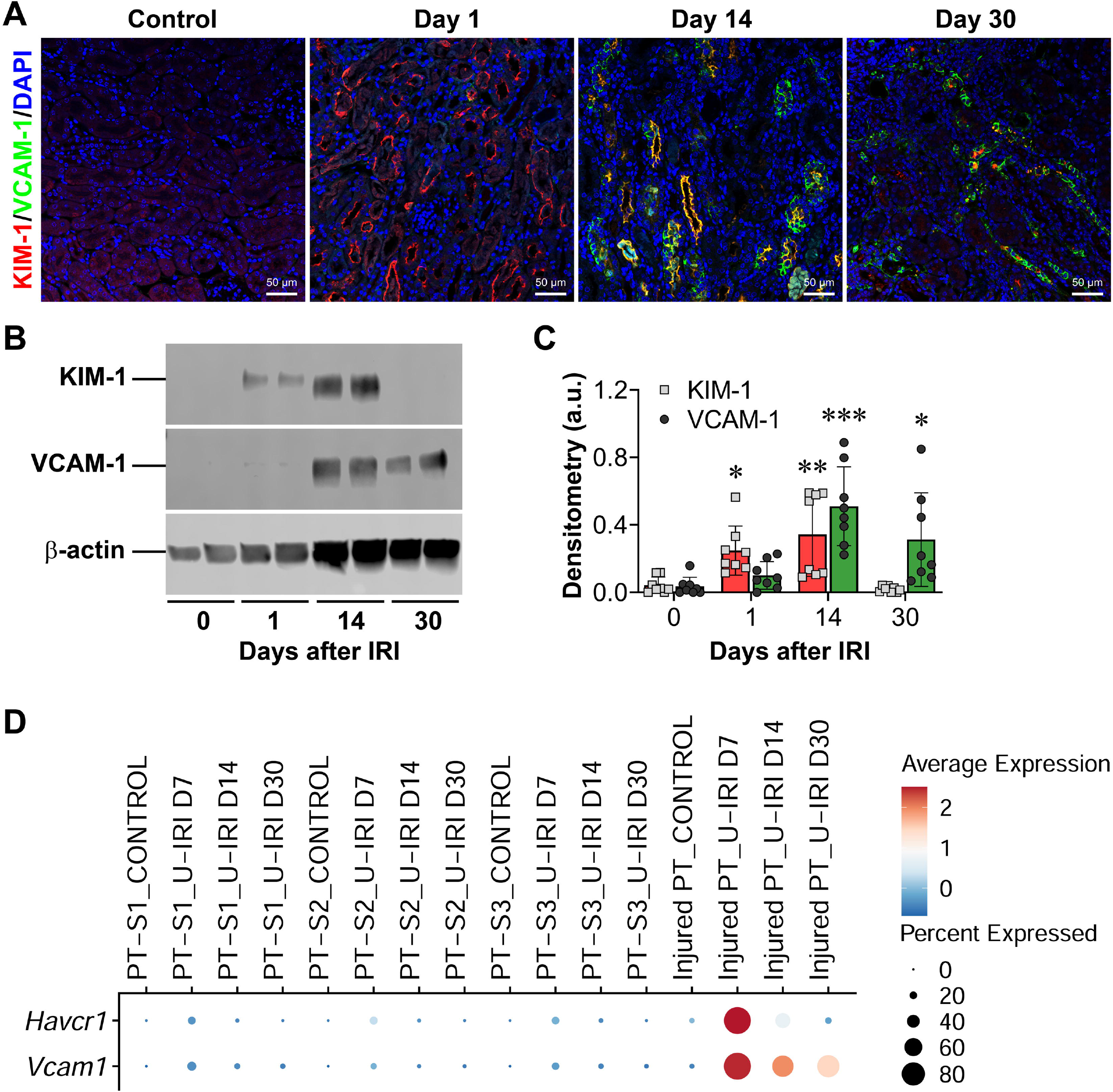
U-IRI leads to an increase of VCAM-1 expression in the proximal tubules during CKD progression. *Wild-type* mice were subjected to unilateral ischemia/reperfusion injury (U-IRI) for 27 minutes and sacrificed on day 1, 14 and 30 after injury. (A) Kidney sections were immunostained with KIM-1 (green) and VCAM-1 (red) and imaged on confocal microscope. Scale bars, 50 μm. (B) Western blot analysis for the protein expression of KIM-1, VCAM-1, and β-actin (re-probed on the same blot after stripping) was performed on whole kidney lysates (each lane is from a separate kidney) on uninjured control kidneys and injured kidneys 1, 14, and 30 days after IRI. The uncropped scans of all blots were supplied in the Supplementary Figure 1. (C) Densitometry was analyzed using ImageJ. Both KIM-1 and VCAM-1 protein expression levels were normalized to β-actin protein expression levels. Data are presented as mean ± SD. n=8 kidneys/group. p<0.001 (KIM-1) and p<0.0001 (VCAM-1) by one-way ANOVA. *p<0.05, **p<0.01, and ***p<0.001 as compared to day 0 by Tukey multiple comparison. (D) The distribution and relative gene expression of *Havcr1* and *Vcam1* across proximal tubule S1, S2, S3, and injured proximal tubule cell clusters in the control kidneys and injured kidneys 1, 14, and 30 days after U-IRI were visualized in a dot plot. Abbreviation: PT, proximal tubule; D, day.

**Table 3.**
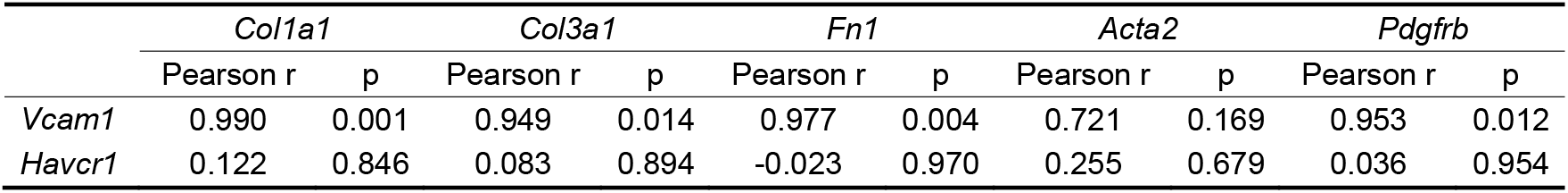
Pearson correlation coefficient with two-tailed p value analysis of gene expression.

### Inflammation milieu promotes proximal tubule VCAM-1 expression

It has been well documented that proinflammatory cytokines can induce endothelial cells to express VCAM-1, which in turn mediates firm adhesion and spreading of leukocytes on the endothelial cell surface (23). A recent single-nucleus assay for transposase accessible chromatin sequencing (snATAC-seq) analysis of adult human kidney reveals that NF-κB regulates the molecular signature of a subpopulation of proximal tubules that express VCAM1 (24). Morphologically, the lumen of some injured proximal tubules is filled with casts (25). At the single cell transcriptome level, we found that proinflammatory cytokines including *Tnf, Il1b*, and *Ifng* were highly upregulated by the infiltrated immune cells. To identify the signals that induce proximal tubule VCAM-1 expression, we treated MPT cells with various stimuli that were present at the late stage of IRI and found that neither serum starvation, exposure to cell debris that mimic cast materials, oxidative stress induced by H_2_O_2_, or cytokine interferon γ could induce *Vcam1* expression in the immortalized proximal tubule cells (Figure 2A). However, septic shock induced by LPS and inflammation induced by TNFα significantly induced *Vcam1* expression by 12- and 11-fold, respectively, in MPT cells as compared to the PBS control treatment (Figure 2A). In contrast, LPS and TNFα only induced *Havcr1* expression by 2- and 3-fold, respectively (Figure 2A). These data suggest that activation of TLR4/Myd88/NF-κB and TNFR1/TRAF2/NF-κB predominantly induce *Vcam1* but not *Havcr1* expression in the proximal tubule epithelial cells. Because IRI induces sterile inflammation but not sepsis, we primarily focus on the proinflammatory cytokines in the following studies.

**Figure 2.**
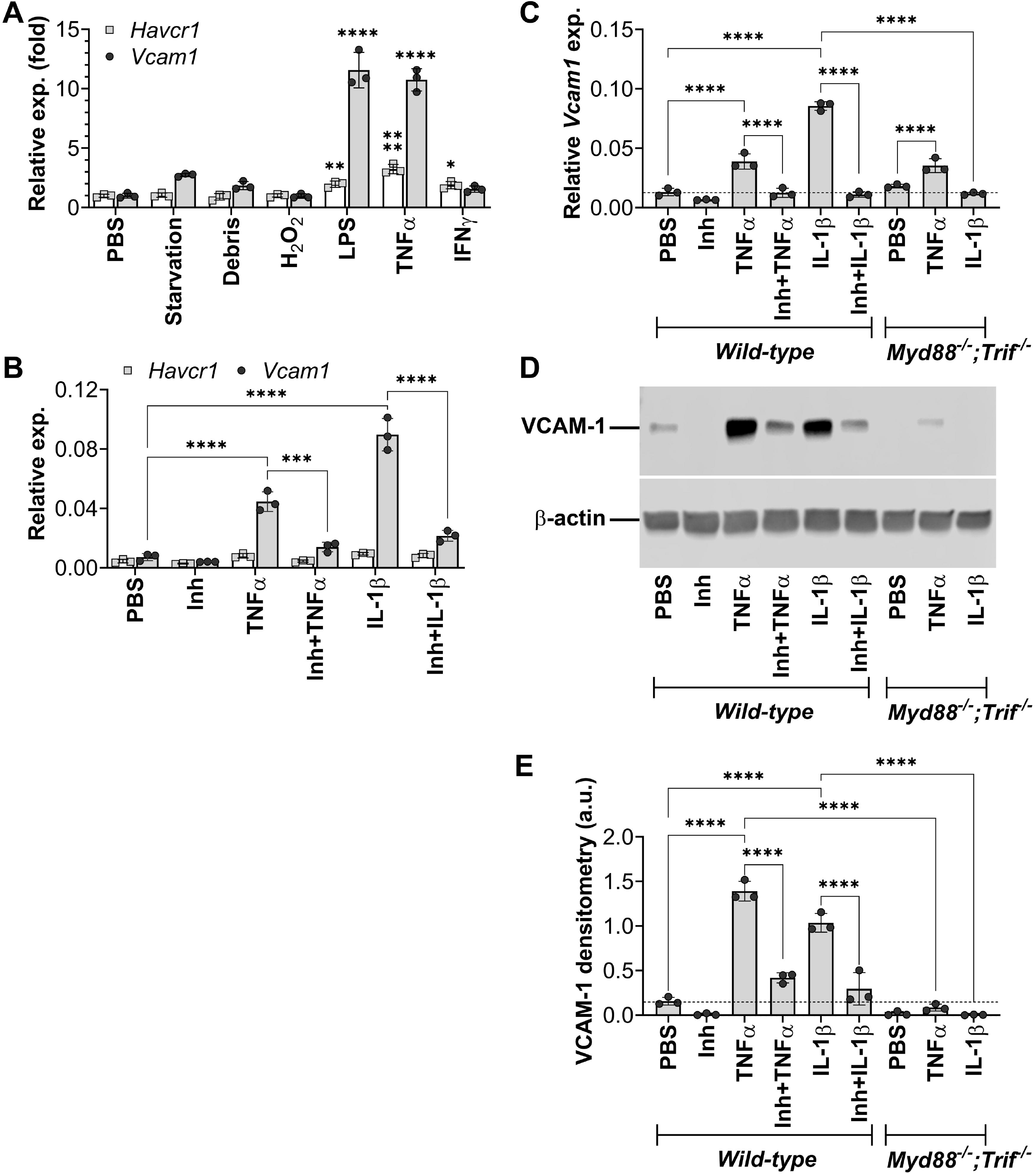
Proinflammatory cytokines promote VCAM-1 expression in proximal tubular epithelial cells. (A) MPT cells were treated with PBS (control), serum starvation, MPT cell debris, H_2_O_2_ (500 μM), LPS (10 ng/mL), TNFα (20 ng/mL), or interferon γ (100 ng/mL) for 6 hours. Quantitative PCR for *Havcr1* and *Vcam1* on MPT mRNA. p<0.0001 by one-way ANOVA. n=3/condition. *p<0.05, **p<0.01 and ****p<0.0001 as compared to PBS by Tukey multiple comparison. (B) MPT cells were treated with PBS (control), NF-κB inhibitor (1μM), TNFα (20 ng/mL) ± NF-κB inhibitor (1 μM), or IL-1 β ± NF-κB inhibitor (1 μM) for 6 hrs. Quantitative PCR for *Havcr1* and *Vcam1* on MPT mRNA. p<0.0001 by one-way ANOVA. n=3/condition. ****p<0.0001 by Tukey multiple comparison. (C-E) *Wild-type* primary cultured renal cells (PCRCs) were treated with PBS (control), NF-κB inhibitor (1μM), TNFα (20 ng/mL) ± NF-κB inhibitor (1 μM), or IL-1β ± NF-κB inhibitor (1 μM). *Myd88^−/−^;Trif^−/−^* PCRCs were treated with PBS (control), TNFα (20 ng/mL), or IL-1β. (C) Quantitative PCR for *Vcam1* on PCRC mRNA after 6 hrs treatment. p<0.0001 by one-way ANOVA. n=3/condition. ****p<0.0001 by Tukey multiple comparison. (D) Western blot analysis for the protein expression of VCAM-1 and β-actin (re-probed on the same blot after stripping) was performed on PCRC protein lysates after 24 hrs treatment. The uncropped scans of all blots were supplied in the Supplementary Figure 2. (E) Densitometry was analyzed using ImageJ. VCAM-1 protein expression levels were normalized to β-actin protein expression levels. Data are presented as mean ± SD. n=3/condition. p<0.0001 by one-way ANOVA. ****p<0.0001 by Tukey multiple comparison.

To test our hypothesis, we treated MPT cells with TNFα and IL-1β in the absence or presence of selective IKKα and IKKβ inhibitor, ACHP (NF-κB inhibitor) and found that as compared to the control PBS treatment, both TNFα and IL-1β markedly induced MPT cells to express *Vcam1*, which was significantly suppressed by co-treatment with NF-κB inhibitor by 68% and 76%, respectively (Figure 2B). Furthermore, we treated primary wild-type cultured renal cells (PCRCs), which contains 72% proximal tubular epithelial cells (10) with TNFα and IL-1β in the absence or presence of NF-κB inhibitor and found that NF-κB inhibitor could abolish VCAM-1 mRNA and protein expression induced by TNFα and IL-1β (Figure 2C-E and Supplementary Figure 2). Double knockout of *Myd88* and *Trif* that blocks IL-1 R/Myd88/NF-κB signaling completely abolished VCAM-1 expression induced by IL-1β but not TNFα (Figure 2C-E and Supplementary Figure 2). Together, these results suggest that VCAM-1 expression is dependent on proinflammatory cytokine-mediated NF-κB activation in the proximal tubular epithelial cells.

### Knockout of *Ccr2* attenuates VCAM-1 expression following U-IRI

We have previously reported that following U-IRI, knockout of Ccr2 decreases the number of macrophages, dendritic cells, and T cells in the tubulointerstitium, which attenuates interstitial fibrosis and reduces expression levels of proinflammatory cytokines including *Tnf* and *Il1b* (12). Here, we used a Proteome Profiler Mouse XL Cytokine Array Kit to systematically detect specific inflammation-related proteins and compared their expression levels between *wild-type* and *Ccr2^−/−^* kidneys. Of 111 different cytokines printed on the membranes, 30 cytokines were detected in the injured kidneys 30 days after U-IRI (Figure 3A and Supplementary Figure 3). Among them, CXCL9, coagulation factor III (CD142, gene symbol *F3*), and VCAM-1 were the top 3 cytokines, of which the expression levels were decrease by 79%, 54%, and 52%, respectively, in the injured *Ccr2^−/−^* kidneys as compared to the *wild-type* kidneys (Figure 3A). As compared to the other known detected kidney injury markers, KIM-1 and ICAM-1, the expression level of VCAM-1 decreased the most in the injured *Ccr2^−/−^* kidneys as compared to the *wild-type* kidneys (Figure 3B). To validate this finding, we analyzed both mRNA and protein levels of VCAM-1 and found that VCAM-1 expression levels were significantly decreased in the injured *Ccr2^−/−^* kidneys as compared to the *wild-type* kidneys (Figure 3C-E and Supplementary Figure 4).

**Figure 3.**
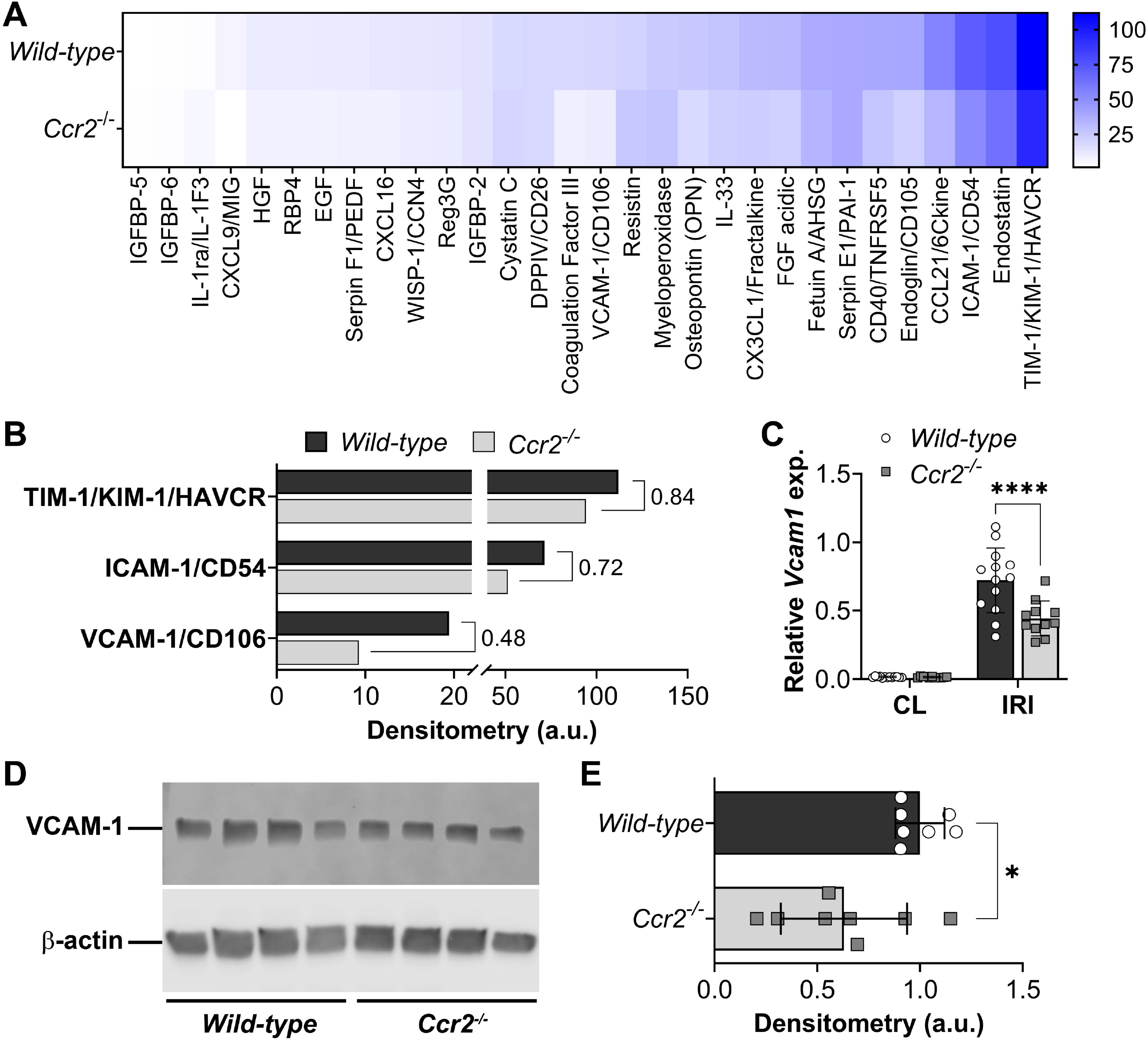
Knockout of *Ccr2* attenuates VCAM-1 expression at the late stage of U-IRI. *Wild-type* and *Ccr2^−/−^* mice were subjected to unilateral ischemia/reperfusion injury (U-IRI) and sacrificed 30 days after injury. (A) Three kidney protein lysates (60 μg) per genotype were pooled and analyzed using the Proteome Profiler Mouse XL Cytokine Array Kit. The array membranes were supplied in the Supplementary Figure 3. Of 111 mouse cytokine proteins targeted by the kit, 30 proteins were detected in a two-minute exposure. The relative densitometry was shown in a heatmap. (B) The densitometry of detected proximal tubular injury markers, TIM-1, ICAM-1 and VCAM-1 were analyzed. (C) Quantitative PCR for *Vcam1* was performed on whole kidney mRNA. n=13 (*Wild-type*) and n=11 (*Ccr2^−/−^*). p<0.0001 by one-way ANOVA. ****p<0.0001 by Tukey multiple comparison. CL, contralateral. (D) Western blot analysis for the protein expression of VCAM-1 and β-actin (re-probed on the same blot after stripping) was performed on whole kidney lysates (each lane is from a separate kidney) on injured *wild-type* and *Ccr2^−/−^* kidneys. The uncropped scans of all blots were supplied in the Supplementary Figure 4. (C) Densitometry was analyzed using ImageJ. VCAM-1 protein expression levels were normalized to β-actin protein expression levels. Data are presented as mean ± SD. n=8 kidneys/genotype. p<0.05 (VCAM-1) by two-tailed t test.

### Single-cell RNA sequencing reveals a distinct *VCAM1*-expressing injured proximal tubule epithelial population in human CKD

To test if our findings in the mouse models are clinically relevant, we performed scRNA-seq analysis of 58,357 cells from 10 participants with CKD and 8 healthy participants from the publicly available KPMP Central Biorepository (Tables 1–2). Using unsupervised clustering, we identified clusters of all major kidney, stromal and immune cell types in the healthy participants and participants with CKD (Figure 4A-B and Supplementary Figure 5A). As compared to healthy reference kidneys, a significant decrease of proximal tubules and an increase of immune cells including PMNs, T cells, and NKT cells were observed in the CKD kidneys (Supplementary Figure 5B-C). When subgrouping participants with CKD by their baseline eGFR (stage 1 as eGFR > 90 mL/min, n=1; stage 2 as eGFR = 60-89 mL/min, n=1; stage 3 as eGFR = 30-59 mL/min, n=7; and stage 4 as eGFR = 15-29 mL/min, n=1), we found that the decrease of proximal tubules and the increase of injured proximal tubules were associated with the stages of CKD progression (Figure 4C-D). Because of limited participants in CKD stages 1, 2, and 4, we compared the cell populations between healthy participants with the participants with stage 3 CKD and found that there was a significant decrease in healthy proximal tubule population but a significant increase of injured and dedifferentiated proximal tubules and immune cells (macrophages, PMNs, T cells, and NKT cells) in the CKD kidneys as compared to the healthy reference kidneys (Supplementary Figure 6). Within the injured proximal tubules, VCAM1 but not HAVCR1 was highly expressed (Figure 4B,C,E), which is consistent with our findings in mouse.

**Figure 4.**
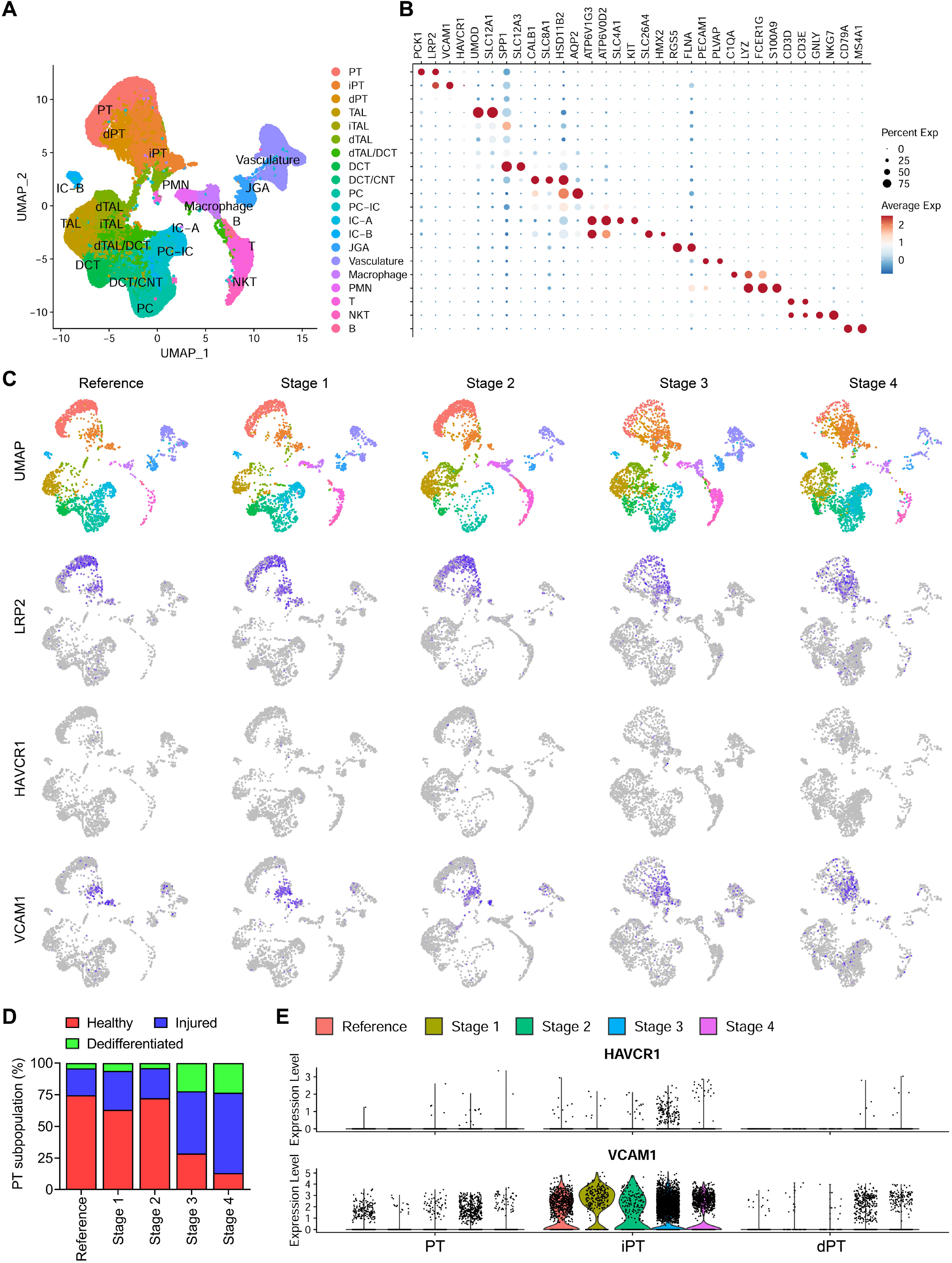
Single-cell transcriptome analysis of patients with CKD identified a distinct injured proximal tubule cell cluster that expresses *VCAM1*. Uniform manifold approximation and projection (UMAP) of 58,357 cells from 10 patients with CKD and 8 healthy participants obtained from the publicly available KPMP Central Biorepository (Tables 1–2; https://atlas.kpmp.org/). Cell clusters were identified using the integrated data from all cells by kidney cell and immune cell lineage-specific marker expression as shown in (B). PT, proximal tubule; iPT, injured PT; dPT, dedifferentiated PT; TAL, thick ascending limb; iTAL, injured TAL; dTAL, dedifferentiated TAL; DCT, distal convoluted tubule; CNT, connecting tubule; PC, principal cell; IC, intercalated cell; JGA, juxtaglomerular apparatus; PMN, polymorphonuclear neutrophil; T, T cell; NKT, natural killer T cell; B, B cell. (C) The integrated data were then split into five sub-datasets: Reference, Stage 1 (eGFR > 90 mL/min), Stage 2 (eGFR = 60-89 mL/min), Stage 3 (eGFR = 30-59 mL/min), and Stage 4 (eGFR = 15-29) based on their baseline eGFR (Table 1). Cell clusters and gene expression level of *LRP2, HAVCR1*, and *VCAM1* of each dataset were visualized in UMAP and feature plots. (D) The percentage of the sub-populations (healthy, injured, and dedifferentiated) of proximal tubule (PT) is provided for each dataset. (E) The distribution and relative gene expression of *HAVCR1* and *VCAM1* in the proximal tubule (PT), injured proximal tubule (iPT), and dedifferentiated proximal tubule (dPT) cell clusters across all the datasets were visualized in the violin plots.

## DISCUSSION

In this work, we found that in response to IRI, the injured kidney markedly upregulated KIM-1 expression, which peaked on day 1 and gradually decreased close to baseline on day 30. In contrast, VCAM-1 was not induced initially on day 1 but markedly increased afterwards and remained highly expressed on day 30 following IRI, which correlates with the progression of kidney atrophy and kidney fibrosis (Table 3). The expression kinetics of these two injury markers is in line with the data reported by Humphrey lab (on day 2, 14, and 42 following bilateral IRI) (22). Histologically, both KIM-1 and VCAM-1 were predominantly expressed by proximal tubular epithelial cells (Figure 1) (22), suggesting that acute induction of KIM-1 and delayed induction of VCAM-1 in proximal tubules demarcates two distinct types of cellular response following AKI and the onset of VCAM-1 might mark the earliest molecular event from AKI-to-CKD transition. A recent genetic lineage tracing study revealed that KIM-1 is induced in the injured proximal tubular cells very early after bilateral IRI, nearly 60% of which co-expressed KI-67 on day 2 after IRI and majority of which account for tubular repair driven by the transcription factor *Foxm1* (17). During AKI-to-CKD transition, as initial injury is preceded, a secondary injury that is mediated by a second wave of immune activation occurs, resulting in a proinflammatory milieu including TNFα and IL-1β (15). These proinflammatory cytokines in turn induced proximal tubular cells to express VCAM-1 but not KIM-1 via NF-κB activation (Figure 2) (26). Here, we showed that blockade of TNFα and IL-1β signaling pathway using NF-κB inhibitor or double knockout of IL-1β downstream signaling mediators *Myd88* and *Trif* markedly suppressed VCAM-1 expression induced by TNFα or IL-1β in vitro (Figure 2) and attenuation of proinflammatory milieu by suppressing recruitment of immune cells using *Ccr2^−/−^* mouse line significantly decreases kidney VCAM-1 expression 30 days following U-IRI in vivo (Figure 3). Consistent with our findings from in vitro and in vivo models, the single cell transcriptome analysis of patients with CKD revealed a distinct cell cluster of injured proximal tubular cells that highly expressed *VCAM1* but not *KIM1* (Figure 4).

It has been well documented that VCAM-1 functions as an adhesion molecule of lymphocytes, monocytes, eosinophils, and basophils to vascular endothelium. In the early 90s, a study of 49 biopsies from patients with interstitial nephritis, systemic vasculitis with crescentic nephritis, minimal change, IgA, lupus nephropathy, diabetic nephropathy, amyloid, or gout showed a marked expression of VCAM-1 interestingly in the proximal tubules but not vascular endothelial cells (27). With advance of single cell RNA-sequencing technology, a distinct cell cluster of proximal tubules that expressed *Vcam1* were proinflammatory, downregulated expression of terminal differentiation markers and anti-oxidative stress genes, upregulated expression of multiple major histocompatibility complex (MHC) class II genes (*H2-Aa, H2-Ab1, H2-Eb1*, and *Cd74*) and class I genes (*H2-D1* and *H2-K1*), and became “failed repair proximal tubule cells (FR-PTCs)” after bilateral IRI (15, 22). This cluster of FR-PTCs could be also detected in a folic acid nephropathy mouse model and over time in human kidney allografts (22, 28, 29). In the patients with systemic lupus erythematosus (SLE), urinary VCAM-1 levels were significantly higher than in the healthy controls, and it was closely associated with the severity of renal insufficiency (30). Together, these data suggest a potential crosstalk between proximal tubule cells (but not endothelial cells) and immune cells, which may promote sustained proximal tubule injury during AKI-to-CKD transition.

Mechanistically, blockage of VCAM-1 using neutralizing antibodies against VCAM-1 or genetic ablation of *Vcam1* has been shown to reduce disease severity and inflammation in various mouse models including brain aging (microglial reactivity and cognitive deficits) (31), atopic dermatitis (32), nonalcoholic steatohepatitis (NASH) (33), angiotensin II-induced hypertension (34), etc. In kidney, one study demonstrated that endothelium-specific knockout of hypoxia-inducible transcription factor 2α (*Hif2a*) led to an increased expression of renal injury markers and inflammatory cell infiltration in the injured kidney 3 days after U-IRI, which could be reversed by blockade of VCAM-1 and very late antigen-4 (VLA4) using monoclonal antibodies (35). Interestingly, blockade of VCAM-1 and VLA4 didn’t protect kidney from injury 3 days after U-IRI in the control mice, suggesting that dysregulation of VCAM-1 expression, most likely in the endothelium, is HIF-2α-dependent during early phase of IRI. However, the function of VCAM-1 in the proximal tubules during AKI-to-CKD transition remains poorly understood, and additional studies are needed to investigate whether proximal tubularVCAM-1 functions more than an injury marker during CKD progression.

## Acknowledgements

We are grateful to Dr. Ruslan Medzhitov from the Department of Immunobiology at the Yale School of Medicine for the generous gift of *Myd88^−/−^:Trif^−/−^* mice.

## Author Contributions

IM and KG performed the primary experiments, analyzed the data, and edited the manuscript. JG prepared PCRCs, interpreted the data, and edited the manuscript. LX designed the experiments, performed the scRNA-seq analyses, oversaw the project, and wrote the manuscript.

## Data Sharing Statement

All data are available within the manuscript/supporting information or from the authors upon request.

## Supplemental Table of Contents

Supplementary Figure 1. U-IRI leads to an increase of VCAM-1 expression in the kidney during CKD progression.

Supplementary Figure 2. Proinflammatory cytokines promote VCAM-1 expression in proximal tubular epithelial cells.

Supplementary Figure 3. Knockout of Ccr2 attenuates VCAM-1 expression

Supplementary Figure 4. Knockout of Ccr2 attenuates VCAM-1 expression at the late stage of U-IRI.

Supplementary Figure 5. Single-cell transcriptome analysis of patients with CKD and healthy participants.

Supplementary Figure 6. The percentage of different cell populations.

