## Supplementary Information for "Inflammation-mediated Upregulation of VCAM-1 but not KIM-1 during Acute Kidney Injury to Chronic Kidney Disease Transition"

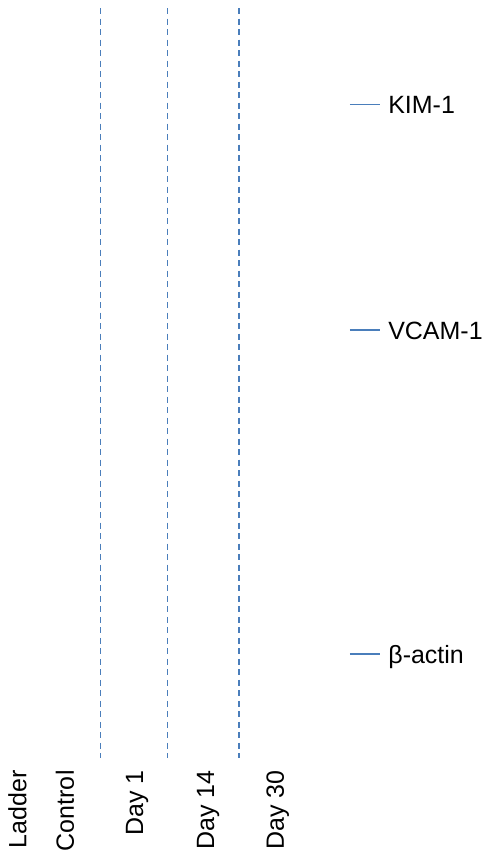

**Supplementary Figure 1. U-IRI leads to an increase of VCAM-1 expression in the kidney during CKD progression.** *Wild-type* mice were subjected to 27 minutes of unilateral ischemia/reperfusion injury (U-IRI) and sacrificed on day 1, 14 and 30 after injury. Western blot analysis for the protein expression of KIM-1, VCAM-1, and β-actin (re-probed on the same blot after stripping) was performed on whole kidney lysates (each lane is from a separate kidney) on uninjured control kidneys and injured kidneys 1, 14, and 30 days after IRI.

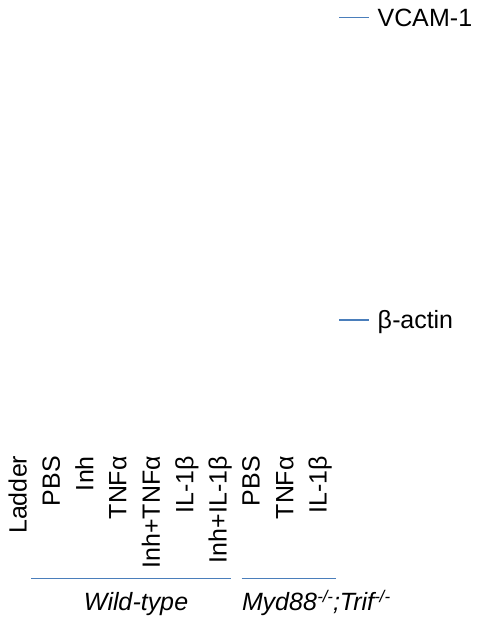

**Supplementary Figure 2. Proinflammatory cytokines promote VCAM-1 expression in proximal tubular epithelial cells.** *Wild-type* primary cultured renal cells (PCRCs) were treated with PBS (control), NF-κB inhibitor (1 µM), TNFα (20 ng/mL) ± NF-κB inhibitor (1 µM), or IL-1β ± NF-κB inhibitor (1 µM). *Myd88^-/-^;Trif^-/-^* PCRCs were treated with PBS (control), TNFα (20 ng/mL), or IL-1β (20 ng/mL). Western blot analysis for the protein expression of VCAM-1 and β-actin (re-probed on the same blot after stripping) was performed on PCRC protein lysates after 24 hrs treatment.

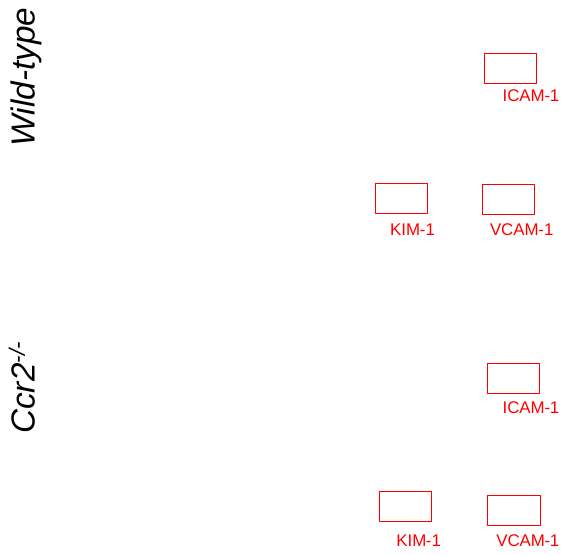

**Supplementary Figure 3. Knockout of *Ccr2* attenuates VCAM-1 expression at the late stage of U-IRI.** *Wild-type* and *Ccr2^-/-^* mice were subjected to 27 minutes of unilateral ischemia/reperfusion injury (U-IRI) and sacrificed 30 days after injury. Three kidney protein lysates (60 μg) per genotype were pooled and analyzed using the Proteome Profiler Mouse XL Cytokine Array Kit. The array membranes were exposed to X-ray film for 2 min at the same time.

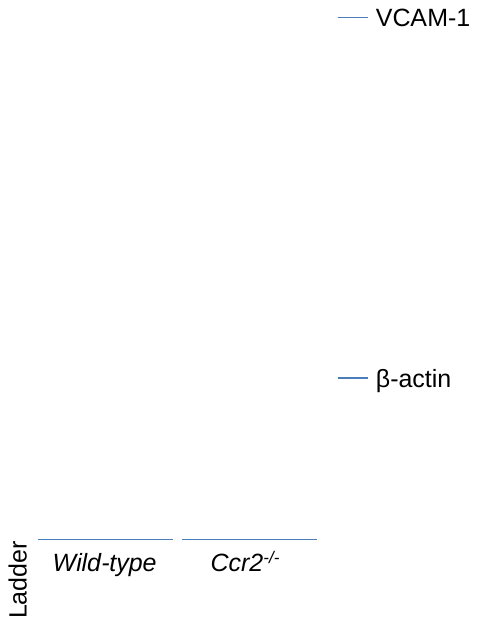

**Supplementary Figure 4. Knockout of *Ccr2* attenuates VCAM-1 expression at the late stage of U-IRI.** *Wild-type* and *Ccr2^-/-^* mice were subjected to 27 minutes of unilateral ischemia/reperfusion injury (U-IRI) and sacrificed 30 days after injury. Western blot analysis for the protein expression of VCAM-1 and β-actin (re-probed on the same blot after stripping) was performed on whole kidney lysates (each lane is from a separate kidney) on injured *wild-type* and *Ccr2^-/-^* kidneys.

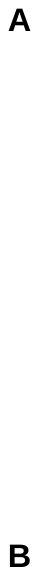

**Supplementary Figure 5. Single-cell transcriptome analysis of patients with CKD and healthy participants.** (A) Uniform manifold approximation and projection (UMAP) of 58,357 cells from 10 patients with CKD (44,782 cells) and 8 healthy participants (13575 cells) obtained from the publicly available KPMP Central Biorepository (Tables 1-2). (B) The proportions of the cell populations are provided for each group.

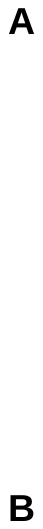

**Supplementary Figure 6. The percentage of different cell populations.** (A) The percentages of the indicated cell populations are provided between healthy participants and patients with CKD. n=8 (Reference) and n=10 (CKD). *p<0.05 and **p<0.01 by two-tailed t test. (B) The percentages of the indicated cell populations are provided between healthy participants and patients with Stage 3 CKD. n=8 (Reference) and n=10 (CKD). *p<0.05 and **p<0.01 by two-tailed t test. PT, proximal tubule; iPT, injured PT; dPT, dedifferentiated PT; PMN, polymorphonuclear neutrophil; T, T cell; NKT, natural killer T cell; B, B cell.
